# A functional non-coding RNA is produced from *xbp-1* mRNA

**DOI:** 10.1101/2020.03.20.000869

**Authors:** Xiao Liu, Jean-Denis Beaudoin, Carrie Ann Davison, Sara G. Kosmaczewski, Benjamin I. Meyer, Antonio J. Giraldez, Marc Hammarlund

**Affiliations:** Department of Genetics, Yale University School of Medicine, New Haven, CT, USA; Yale Stem Cell Center, Yale University School of Medicine, New Haven, CT, USA; Department of Neuroscience, Yale University School of Medicine, New Haven, CT, USA

## Abstract

The *xbp-1* mRNA encodes the XBP-1 transcription factor, a critical part of the unfolded protein response. Here we report that an RNA fragment produced from *xbp-1* mRNA cleavage is a biologically active non-coding RNA (ncRNA) in *Caenorhabditis elegans* neurons, providing the first example of ncRNA derived from mRNA cleavage. We show that the *xbp-1* ncRNA is crucial for axon regeneration *in vivo*, and that it acts independently of the protein-coding function of the *xbp-1* transcript. Structural analysis indicates that the function of the *xbp-1* ncRNA depends on a single RNA stem; and this stem forms only in the cleaved *xbp-1* ncRNA fragment. Disruption of this stem abolishes the non-coding but not coding function of the endogenous *xbp-1* transcript. Thus, cleavage of the *xbp-1* mRNA bifurcates it into a coding and a non-coding pathway; modulation of the two pathways may allow neurons to fine-tune their response to injury and other stresses.

**Graphic abstract:** 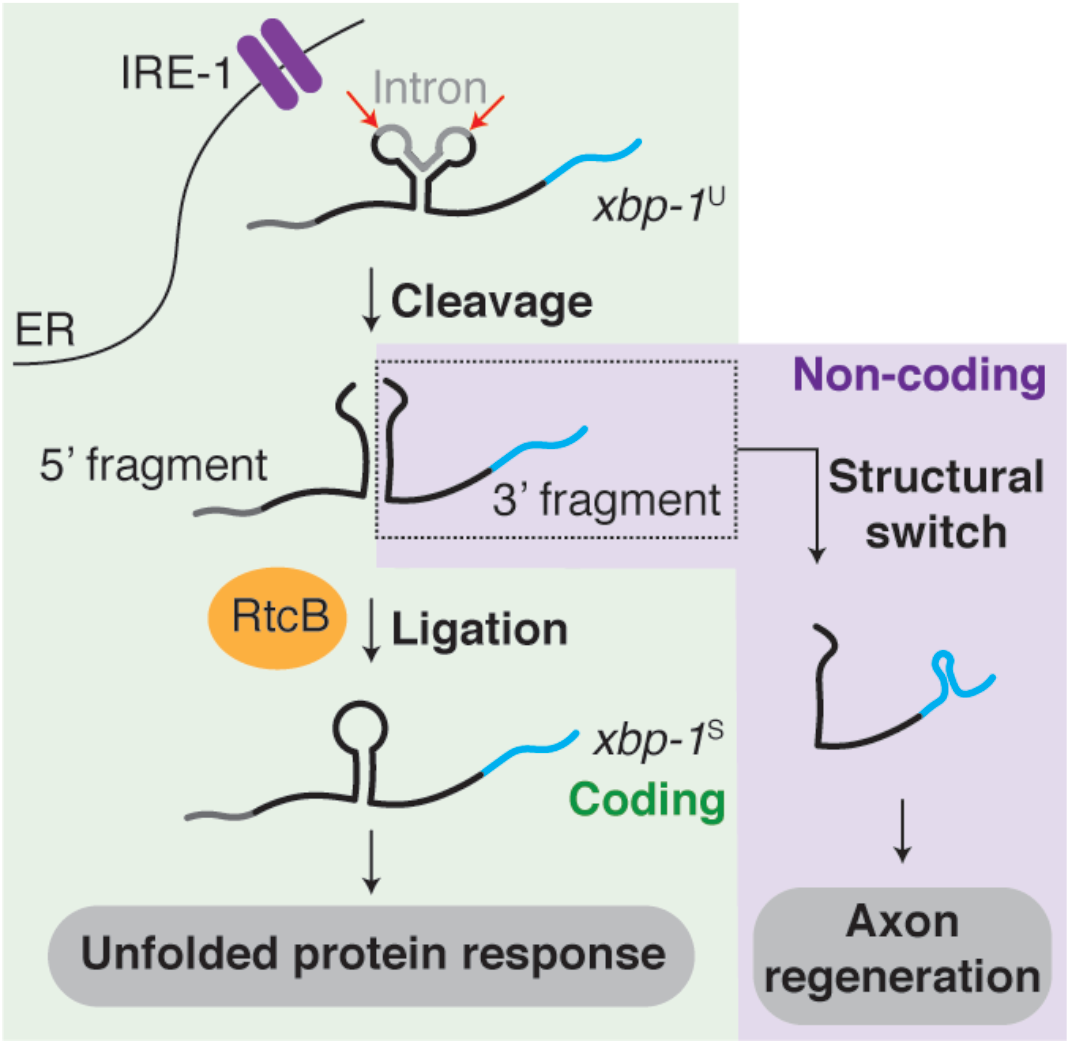

## Introduction

The IRE/XBP branch of the unfolded protein response (XBP-UPR) is conserved from yeast to human and is a critical component of the cellular response to protein stress in the ER (Walter and Ron, 2011). The output of the XBP-UPR is activity of the XBP-1 protein, a DNA-binding transcription factor encoded by the *xbp-1* locus. Translation of active XBP-1 protein requires processing of *xbp-1* mRNA by a non-canonical cytoplasmic RNA splicing event. This splicing event removes a short central sequence from the *xbp-1* mRNA, resulting in a frame shift that brings the functional C-terminal half of the XBP-1 protein into frame. Mechanistically, cytoplasmic *xbp-1*^U^ mRNA is first cleaved twice by the endonuclease IRE-1(Calfon et al., 2002; Kawahara et al., 1998; Sidrauski and Walter, 1997; Yoshida et al., 2001). Next, the 5’ and 3’ fragments are ligated by the RNA ligase RtcB to generate the spliced *xbp-1*^S^ mRNA that encodes the XBP-1 protein (Fig. 1A)(Jurkin et al., 2014; Kosmaczewski et al., 2014; Lu et al., 2014; Sidrauski et al., 1996). The XBP-UPR is crucial for cellular protein homeostasis, and dysregulation of *xbp-1* splicing is implicated in inflammatory diseases, metabolic disease and several types of cancer (Jiang et al., 2015). In the nervous system, the function of the XBP-UPR has been associated with a wide range of neurodegenerative diseases including Alzheimer’s disease, amyotrophic lateral sclerosis (ALS), Huntington’s disease, and Parkinson’s disease (Hetz and Mollereau, 2014; Jiang et al., 2015).

**Fig. 1.**
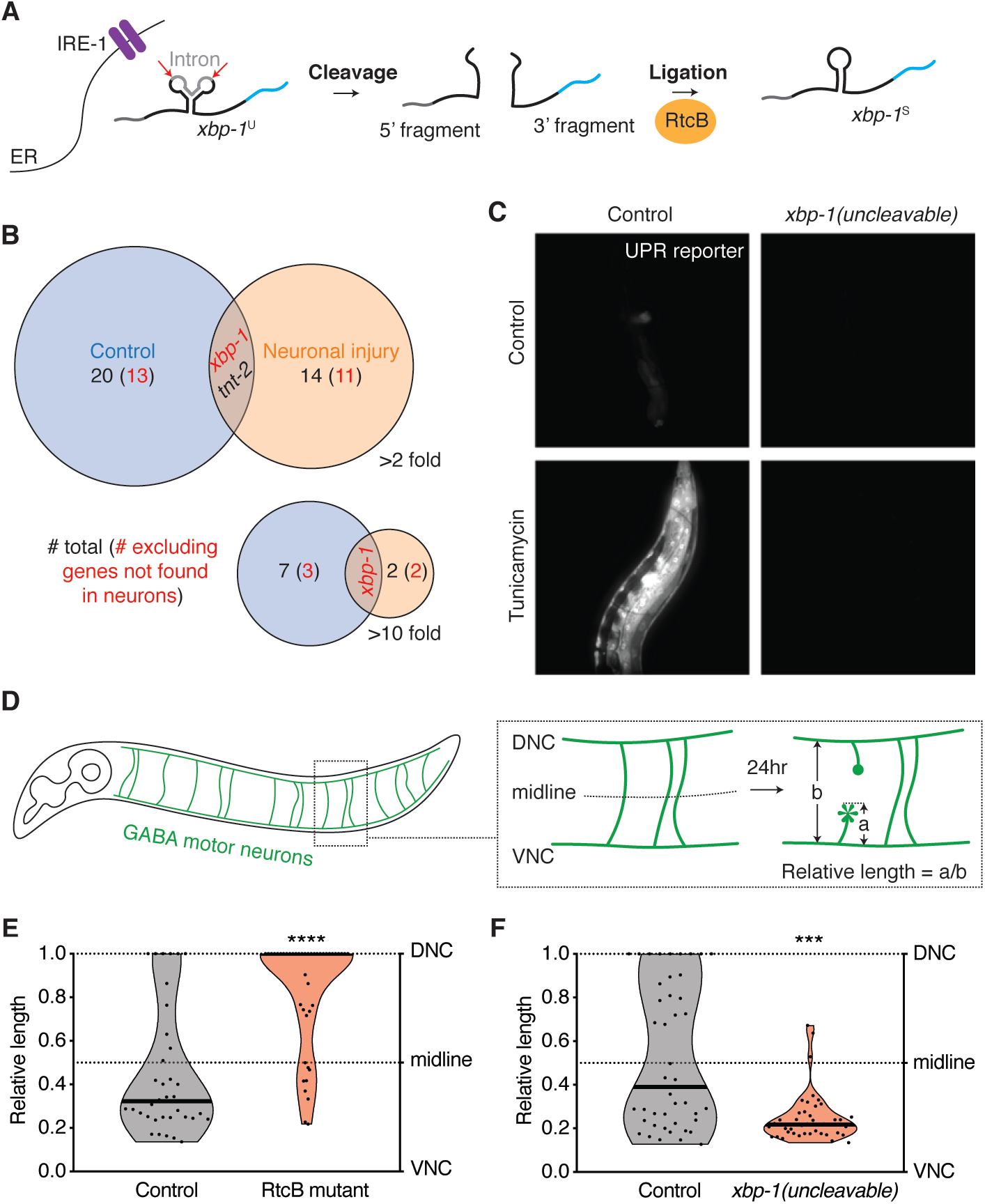
A processing intermediate of the *xbp-1* mRNA promotes axon regeneration. (**A**) Diagram of the *xbp-1* mRNA splicing pathway. (**B**) RNA-seq analysis identifies non-canonical RNA junctions that are enriched in RtcB mutant animals compared to non-mutant controls. The Venn diagram plots genes with such RtcB-dependent junctions identified under normal condition (blue) or with neuronal injury (orange) at the indicated fold-change cutoff. (**C**) Animals with uncleavable *xbp-1* fail to mount the UPR either in control conditions or upon tunicamycin treatment (5μg/ml, 24h). (**D**) Scheme of axotomy in *C. elegans* GABA neurons. (**E**) Animals deficient of the RNA ligase RtcB show significantly higher regeneration. *n* = 37 and 44 from left to right. (**F**) Axon regeneration is eliminated in animals with the *xbp-1(uncleavable)* allele. *n* = 44 and 43 from left to right. In (E) and (F), black bar represents the median. \*\*\**P*<0.001, \*\*\*\**P*<0.0001, 2-tailed K-S test.

We previously found that RtcB, the RNA ligase that is required for *xbp-1* splicing and the XBP-UPR, has a very strong effect on axon regeneration in *C. elegans* neurons (Kosmaczewski et al., 2015). Neurons can respond to axon injury by initiating axon regeneration to restore structure and function. Both an unbiased functional screen (Nix et al., 2014) and detailed genetic analysis (Kosmaczewski et al., 2015) indicated that RtcB mutants have extremely high regeneration, among the strongest effects seen. This result was surprising, since neuronal injury is a form of cellular stress, and RtcB mutants completely lack the XBP-UPR and die quickly when treated with tunicamycin to disrupt ER protein homeostasis (Kosmaczewski et al., 2014). Further experiments showed that the effect of RtcB required its ligase activity, but was independent of the XBP-UPR. It was also independent of tRNA ligation, the other activity of RtcB ligation (Kosmaczewski et al., 2015). These data indicate that RtcB affects axon regeneration by an unknown mechanism involving RNA ligation.

Here we show that the *xbp-1* locus, in addition to encoding the XBP-1 protein, also encodes a biologically active non-coding RNA (ncRNA). The *xbp-1* ncRNA is the 3’ fragment produced by cleavage of the cytoplasmic *xbp-1*^U^ mRNA. The effect of RtcB on axon regeneration is due to modulation of the *xbp-1* ncRNA function, rather than the canonical output of the XBP-1 protein. We show that the function of the *xbp-1* ncRNA is context-dependent, and that it is not functional before cleavage, when it is still part of the *xbp-1*^U^ mRNA. We identify a rearrangement of secondary structure that correlates with *xbp-1* ncRNA activity, and show that a single base pair within this structure is essential for ncRNA function.

ncRNAs are produced from various origins and biogenesis pathways. In addition to being generated from distinct regions from the coding RNAs, ncRNAs can also be transcribed from genomic regions that are closely linked to coding genes. For example, many miRNAs and circRNAs are processed from intronic regions of pre-mRNAs (Fu, 2014). However, it is less clear whether ncRNAs can be produced from functional, mature mRNAs. Only until recently has it been postulated that coding RNAs can also have non-coding functions (Crerar et al., 2019; Kumari, 2015; Sampath and Ephrussi, 2016). By characterizing the non-coding function of the *xbp-1* locus, our study provides the first example of a ncRNA directly derived from the cleavage of a mature mRNA. Discovery of a ncRNA pathway that coexists with the coding function of the *xbp-1* locus opens a new window into the molecular mechanisms neurons use to respond to cellular conditions.

## Results

### Blocking *xbp-1* cleavage versus ligation results in opposite phenotype in axon regeneration

RtcB mutant animals, which lack the XBP-UPR completely because the *xbp-1* RNA fragments cannot be ligated to produce the protein-coding *xbp-1*^S^, have extremely high axon regeneration (Kosmaczewski et al., 2015). However, the high regeneration phenotype in these mutants is not caused by the loss of the XBP-UPR, as either inactivating or activating the XBP-UPR does not phenocopy or rescue RtcB mutants and overall has relatively minor effects on regeneration (Kosmaczewski et al., 2015). The high regeneration phenotype is also not due to lack of tRNA maturation, despite the clear role of RtcB in ligating tRNAs (Englert et al., 2011; Popow et al., 2011; Tanaka and Shuman, 2011).

We considered the possibility that an unidentified RNA substrate mediates the effect of RtcB ligation on axon regeneration. We performed RNA-seq comparing RtcB mutant animals and non-mutant controls, and searched for RtcB-dependent RNA junctions that did not have the canonical sequences used by the spliceosome (see Methods). To account for the possibility that neuron stress was required, we sequenced both groups with and without neuronal injury, using a mutation in β-spectrin (*unc-70*) to trigger spontaneous axon breaks (Hammarlund et al., 2007). The analysis (Table S1) confirmed that RtcB is required for production of the *xbp-1*^S^ mRNA, as the RtcB-dependent junction was >500 fold enriched in presence of RtcB compared to RtcB mutant animals. Further, this enrichment was observed in both normal conditions and in the β-spectrin mutant background, suggesting that *xbp-1* splicing occurs constitutively and is not significantly altered by neuronal injury. However, when comparing all junctions using a foldchange cutoff at 10, we identified only 3 genes when we compared the two groups with neuronal injury, and 8 genes when without. Other than *xbp-1*, there was no overlap between the two sets of genes (Fig. 1B, Table S2). The effect of RtcB on axon regeneration is neuron-autonomous (Kosmaczewski et al., 2015), but many of the genes we identified are not expressed in neurons (Fig. 1B, Table S2). Further, many of these junctions were located in non-coding regions near the 5’ or 3’ ends of transcripts, and likely have little effect on gene function (Table S2). Overall, we did not identify strong candidates other than *xbp-1* for RtcB-mediated ligation.

Thus, we considered the possibility that another function of the *xbp-1* gene, independent of the XBP-UPR, might affect axon regeneration. To better characterize *xbp-1* function, we sought to block *xbp-1*^U^ mRNA processing at the cleavage step, rather than at the ligation step (as in the RtcB mutants). Cleavage of *xbp-1*^U^ mRNA is mediated by the IRE-1 endonuclease. However, IRE-1 has many RNA targets (Hollien and Weissman, 2006; Hollien et al., 2009). Further, unlike *xbp-1* mutants, *ire-1* mutant animals are visibly sick. These data suggest that IRE-1’s additional targets are biologically important, and might confound the study of the XBP-UPR. Therefore, to block specifically the cleavage of *xbp-1*^U^ mRNA, we used CRISPR techniques to generate a novel allele, *xbp-1(uncleavable)* (Fig. S1A). Cleavage of the *xbp-1^U^* mRNA by IRE-1 requires a conserved RNA motif at the two cleavage sites (Gonzalez et al., 1999). We introduced mutations into the endogenous *xbp-1* locus that alter these motifs, but that do not affect amino acid coding in either the *xbp-1*^U^ or the *xbp-1*^S^ frame (Fig. S1B).

Both cleavage and ligation are required for the production of the *xbp-1*^S^ transcript that encodes the XBP-1 protein (Fig. 1A). Consistent with this, *xbp-1(uncleavable)* animals could neither produce *xbp-1*^S^ (Fig. S1C) nor activate the XBP-UPR (Fig. 1C), similar to RtcB ligase mutants (Kosmaczewski et al., 2014). Interestingly, however, we found that these two mutants have completely opposite phenotypes in axon regeneration. The RtcB ligase mutants have extremely high axon regeneration (Fig. 1D, E), while the *xbp-1(uncleavable)* allele eliminates regeneration (Fig. 1F). These data suggest that an *xbp-1* mRNA processing intermediate between the cleavage step and the ligation step functions in axon regeneration.

### The *xbp-1* 3’ fragment promotes axon regeneration independently of the UPR

During *xbp-1*^U^ mRNA processing, IRE-1 cleaves the mRNA twice, generating three fragments: a 5’ fragment, a central fragment, and a 3’ fragment. The 5’ and 3’ fragments are then ligated by RtcB to generate *xbp-1*^S^ mRNA. Thus, blocking cleavage would be expected to result in depletion of all three fragments, while blocking ligation would be expected to cause relative accumulation of the 5’ and 3’ fragments. Thus, the opposite regeneration phenotypes in the cleavage and ligation mutants could be due to a novel function for these intermediate fragments. To determine which intermediate fragment is responsible, we generated *rtcb-1; xbp-1(zc12)* double mutant animals. We found that this allele of *xbp-1*, which contains an early stop codon, suppresses the high regeneration phenotype in RtcB mutant animals (Fig. 2A, ‘control’), likely because nonsense-mediated decay reduces overall *xbp-1* mRNA abundance, affecting all outputs of the *xbp-1* locus. We expressed transgenic *xbp-1* in this double mutant background, with the expectation that expressing the form of *xbp-1* that mediates regeneration would result in the *rtcb-1* phenotype of high regeneration. Expressing *xbp-1* coding sequence along with its native 3’ UTR, either under its native promoter or under a GABA-specific promoter, restored regeneration back to the high *rtcb-1* single mutant level. These data confirm that the function of the *xbp-1* locus in axon regeneration is cell-autonomous, consistent with previous results from RtcB (Kosmaczewski et al., 2015), and indicate that the function of the *xbp-1* locus is mediated by some aspect of the *xbp-1* mRNA. However, when the endogenous *xbp-1* 3’ UTR was swapped with the commonly-used *unc-54* 3’ UTR, expression of *xbp-1* was no longer able to restore regeneration (Fig. 2A). These data indicate that the *xbp-1* 3’ UTR is required to promote axon regeneration in the *rtcb-1; xbp-1(zc12)* context. We concluded that the *xbp-1* 3’ fragment, which contains the 3’ UTR, is likely responsible for the high regeneration phenotype in the RtcB ligase mutant animals. In support of this, we detected with northern blot an accumulation of *xbp-1* 3’ fragments *in vivo* in RtcB mutant animals compared to control (Fig. 2B).

**Fig. 2.**
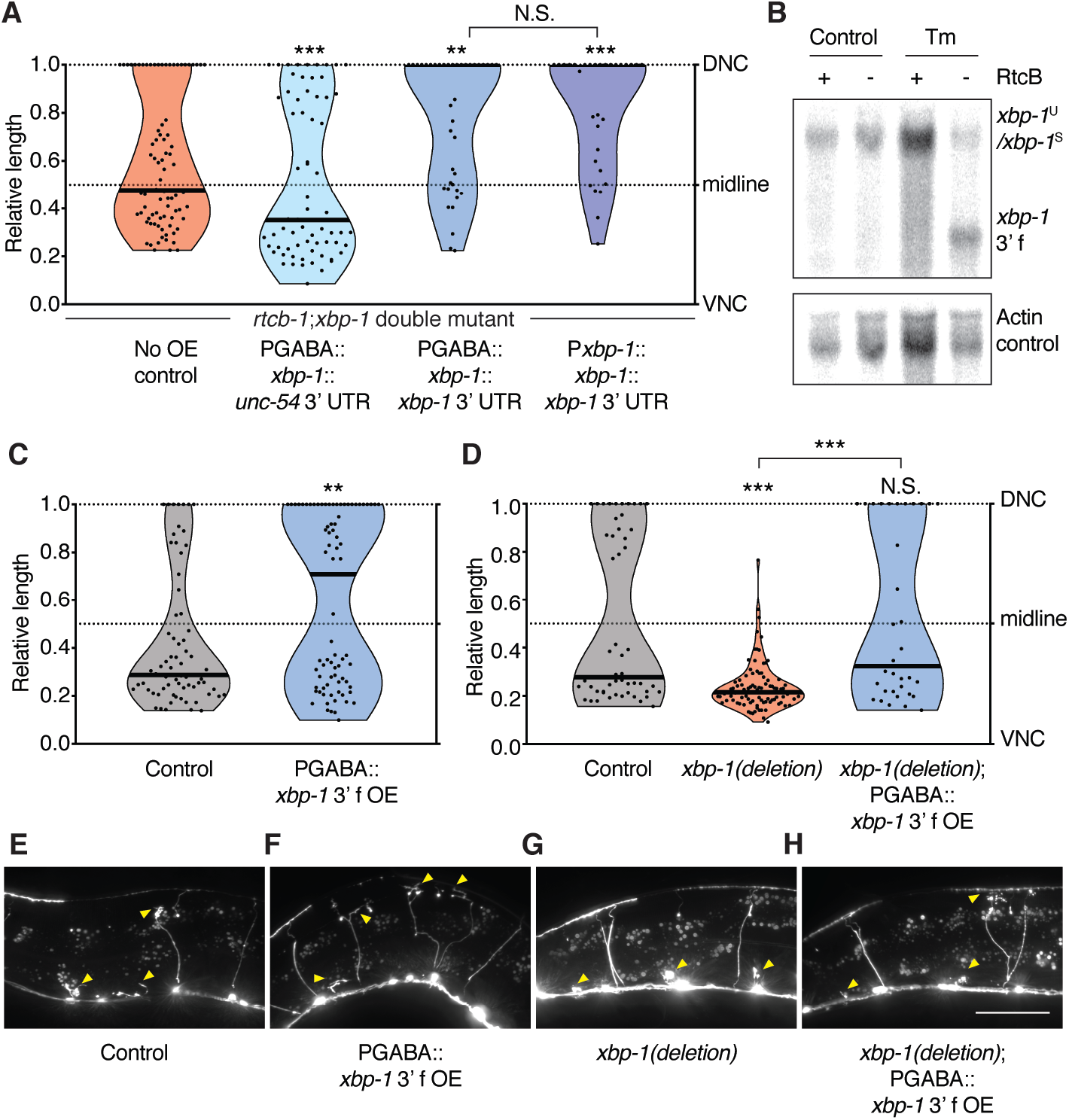
The *xbp-1* 3’ fragment is necessary and sufficient to promote axon regeneration. (**A**) The *xbp-1* 3’ UTR is required to increase regeneration in the *rtcb-1(gk451); xbp-1(zc12)* double mutant background. *n* = 62, 69, 40, and 31 from left to right. (**B**) Northern blot showing *xbp-1* 3’ fragment accumulation in RtcB mutant animals. (**C**) The *xbp-1* 3’ fragment is sufficient to increase regeneration cell-autonomously. *n* = 70 and 79 from left to right. (**D**) Deletion of the genomic *xbp-1* locus eliminates regeneration, which is rescued by GABA-specific expression of the *xbp-1* 3’ fragment. *n* = 52, 92 and 34 from left to right. (**E** to **H**) Representative micrographs of animals of the indicated genotype 24hr post axotomy. Arrows indicate cut axons. Scale bar, 50μm. In (A), (C) and (D), black bar represents the median. N.S., not significant, \*\**P*<0.01, \*\*\**P*<0.001, 2-tailed K-S test.

To further test the function of the *xbp-1* 3’ fragment, and to determine whether it can promote axon regeneration outside the RtcB mutant context, we overexpressed the 3’ fragment specifically in GABA neurons in a non-mutant background. We found that 3’ fragment overexpression increased axon regeneration (Fig. 2C, E, F). To further characterize this effect in relation to the XBP-UPR, we used CRISPR techniques to completely delete the endogenous *xbp-1* gene. These *xbp-1(deletion)* animals had decreased regeneration (Fig. 2D, G), similar to *xbp-1(uncleavable)* (Fig. 1F). In this *xbp-1(deletion)* background, we expressed the *xbp-1* 3’ fragment specifically in GABA neurons. We found that 3’ fragment overexpression restored regeneration to normal levels (Fig. 2D, H). Thus, overexpression of the *xbp-1* 3’ fragment is sufficient to increase regeneration in animals with a normal XBP-UPR and also in animals that completely lack conventional *xbp-1* transcripts and have no XBP-UPR. Further, preventing the production of endogenous *xbp-1* 3’ fragments (in *xbp-1(uncleavable)* (Fig. 1F), and *xbp-1(deletion)* (Fig. 2D)) reduces regeneration below control levels, and this is rescued by the expression of the 3’ fragment (Fig. 2D). These data indicate that the *xbp-1* 3’ fragment is a critical part of the endogenous regeneration response.

### The *xbp-1* 3’ fragment functions as a ncRNA

The *xbp-1* 3’ fragment contains some of the *xbp-1* coding sequence (Fig. S2A), and the overexpression experiments (Fig. 2C, D) included a start codon in the plasmid at the beginning of the *xbp-1* 3’ fragment sequence. We asked whether the *xbp-1* 3’ fragment acts as a non-coding RNA molecule, or whether it encodes a functional peptide. We generated an *xbp-1* 3’ fragment construct with the plasmid start codon frame-shifted (Fig. S2A) and found that this construct retained its ability to promote regeneration (Fig. 3A and 2C), suggesting the coding potential of the *xbp-1* 3’ fragment is dispensable for function in regeneration. Next, we attached only the predicted non-coding (UTR) portion of the *xbp-1* 3’ fragment to the GFP coding sequence, after the GFP stop codon, and expressed this construct (GFP-UTR) specifically in GABA neurons. Expression of this hybrid construct resulted in GFP expression and also increased regeneration. By contrast, expression of GFP with a control 3’ UTR did not affect regeneration, although similar levels of GFP fluorescence were observed (Fig. 3B and S2B). These experiments indicate that the regeneration-promoting activity of the *xbp-1* 3’ fragment is contained within the 3’ UTR, suggesting that the *xbp-1* 3’ fragment acts as a ncRNA.

**Fig. 3.**
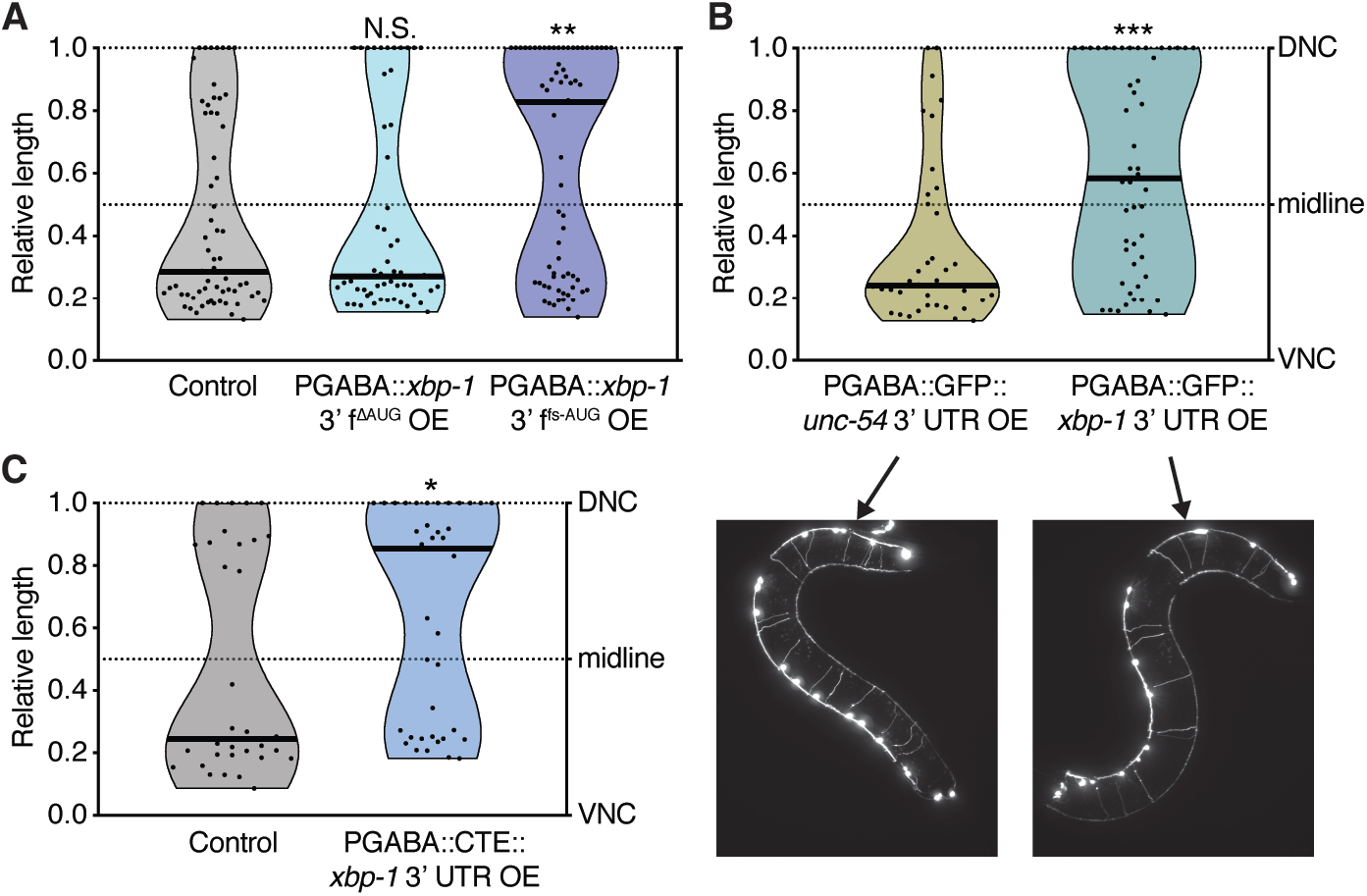
The *xbp-1* 3’ UTR promotes axon regeneration as a ncRNA. **(A)** The *xbp-1* 3’ fragment increases regeneration independently of the identity of the peptide it encodes. ΔAUG, start codon deletion, fs-AUG, frame-shifted start codon. *n* = 63, 55, and 69 from left to right. **(B)** The *xbp-1* 3’ UTR increases regeneration even when fused to GFP coding sequence. Below the violin plot are representative micrographs showing GFP expression. To visualize GFP fluorescence, these constructs were expressed in a wild-type N2 background, as opposed to the *oxIs12* background (GABA-specific GFP marker) used in other axotomy experiments. Regeneration was scored 14h (instead of 24h) after axotomy. *n* = 42 and 50 from left to right. **(C)** When exported from the nucleus into the cytoplasm with the help of CTE, the *xbp-1* 3’ UTR increases regeneration without any coding sequence. *n* = 35 and 39 from left to right. In (A-C), black bar represents the median. N.S., not significant, \**P*<0.05, \*\**P*<0.01, \*\*\**P*<0.001, 2-tailed K-S test.

The endogenous *xbp-1* 3’ fragment is produced in the cytoplasm by cleavage of the *xbp-1*^U^ transcript on the ER membrane. On the other hand, the overexpressed *xbp-1* 3’ fragment is transcribed in the nucleus. Proper pre-mRNA processing and mRNP assembly is required for mRNA stability and efficient export into the cytoplasm (Maniatis and Reed, 2002). Consistent with this idea, we found that the *xbp-1* 3’ fragment only increases regeneration if it contains at least one intron, which is likely to promote nuclear export (Maniatis and Reed, 2002)(Fig. S2C and S2D). Deletion of the start codon (rather than frame-shifting it) also abolished the overexpressed construct’s ability to promote regeneration (Fig. 3A), perhaps due to an effect on mRNA stability or export. To construct a minimal functional sequence without any introns, start codons, or coding potential, we fused the *xbp-1* 3’ UTR with the retrovirus-derived constitutive transport element (CTE). The CTE RNA sequence hijacks the cellular mRNA nuclear export machinery and is exported independently of mRNA processing (Ernst et al., 1997; Grüter et al., 1998). We found that expression of this minimal construct (CTE-UTR) increased axon regeneration (Fig. 3C), similar to expression of the entire *xbp-1* 3’ fragment (Fig. 2C). Together, these data support the idea that the *xbp-1* 3’ fragment acts as a ncRNA to promote regeneration, and that cytoplasmic localization is essential for its function.

### The function of the *xbp-1* ncRNA depends on an RNA stem

To determine which region of the *xbp-1* 3’ fragment is required for its function in regeneration, we tested a series of truncations of the 3’ fragment (Fig. 4A) expressed under a GABA-specific promoter. The results were consistent with our findings that the coding sequence is dispensable, as deletions in this region preserved function (Fig. 4B, Δ1-189 and Δ190-378). By contrast, we found that a 189nt region in the predicted 3’ UTR is required for the 3’ fragment to increase regeneration (Fig. 4B, Δ379-567). *In silico* folding by Vienna RNAfold of this region indicated that it contains a high-probability RNA stem, which was confirmed by SHAPE-MaP (Siegfried et al., 2014) performed on *in vitro* transcribed and folded RNAs (Fig. 4C). We hypothesized that this stem (nucleotides 134-209 from the start of the *xbp-1* 3’ UTR, S^134-209^ hereafter) could be a functional element of the *xbp-1* 3’ fragment. In support of this, deleting S^134-209^ was sufficient to abolish the function of the *xbp-1* 3’ fragment (compare Fig. 4D and 2C).

**Fig. 4.**
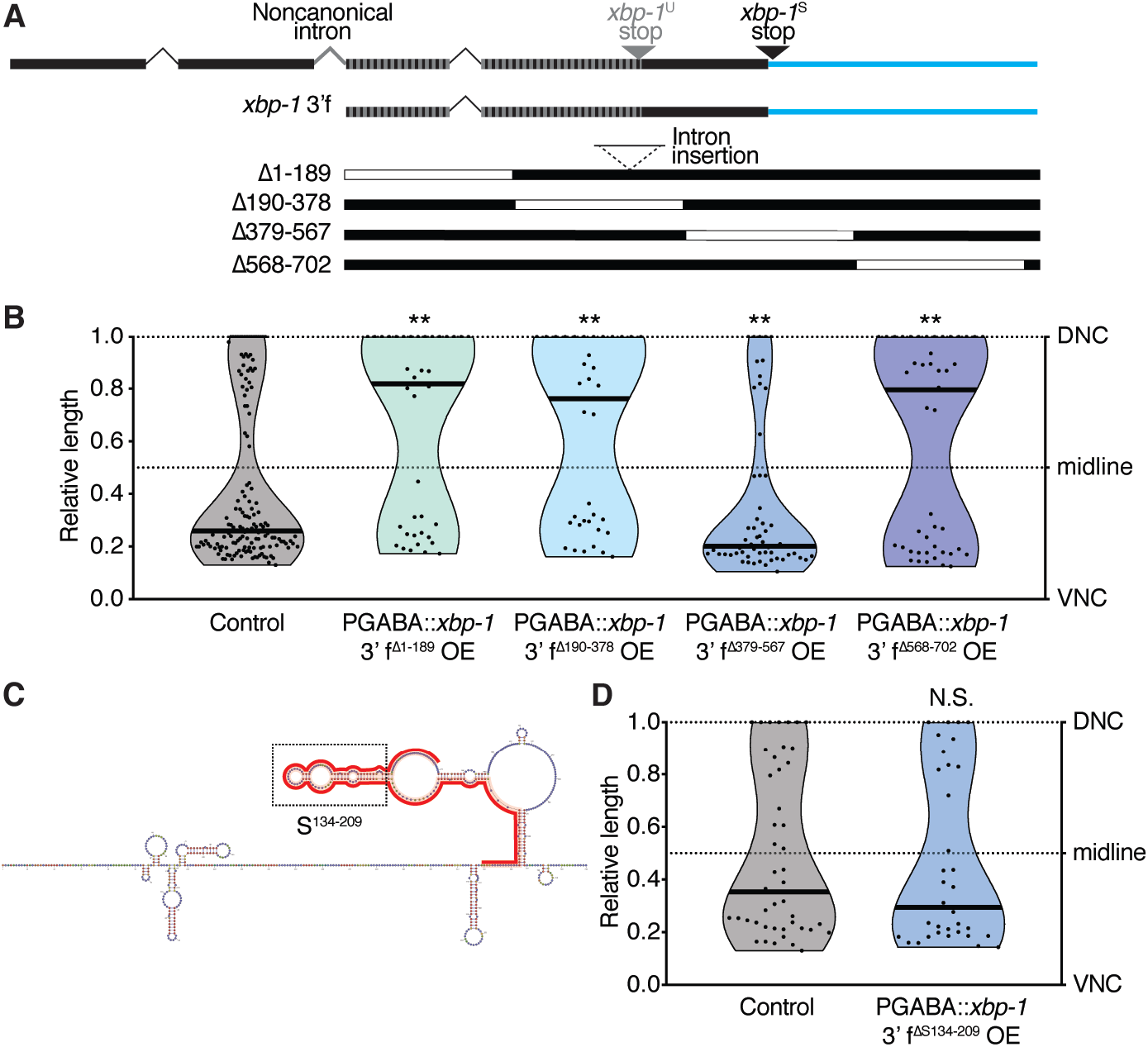
An RNA stem within the *xbp-1* 3’ UTR is required for axon regeneration. (**A**) Diagrams of *xbp-1* 3’ fragment deletion constructs. (**B**) Axotomy results of animals expressing constructs in (A). *n* = 148, 39, 36, 57, and 48 from left to right. (**C**) Predicted secondary structure of the *xbp-1* 3’ fragment based on SHAPE-MaP results. (**D**) The *xbp-1* 3’fragment with the RNA stem S^134-209^ deleted no longer promotes axon regeneration. *n* = 47 and 34 from left to right. In (B) and (D), black bar represents the median. N.S., not significant, \*\**P*<0.01, 2-tailed K-S test.

To further explore the function of S^134–209^, we began by observing that the *xbp-1* 3’ UTR (which contains S^134-209^) does not function in every condition. The *xbp-1* 3’ UTR promotes regeneration when expressed as part of the *xbp-1* 3’ fragment (Fig. 2C), or when attached to the GFP coding sequence (GFP-UTR, Fig. 3B), or when fused to CTE (CTE-UTR, Fig. 3C). By contrast, we found that it does not have a significant effect on regeneration when it is part of the full-length *xbp-1* transcript. The *xbp-1(uncleavable)* allele has poor regeneration even though the 3’ UTR is present (Fig. 1F), and overexpression of either wild-type or uncleavable full-length *xbp-1* with the 3’ UTR under a GABA-specific promoter does not increase regeneration (Fig. 5A). These data suggest that the *xbp-1* 3’ UTR, and perhaps the key S^134-209^ region, is in an inactive structural conformation when part of the full-length *xbp-1* transcript.

**Fig. 5.**
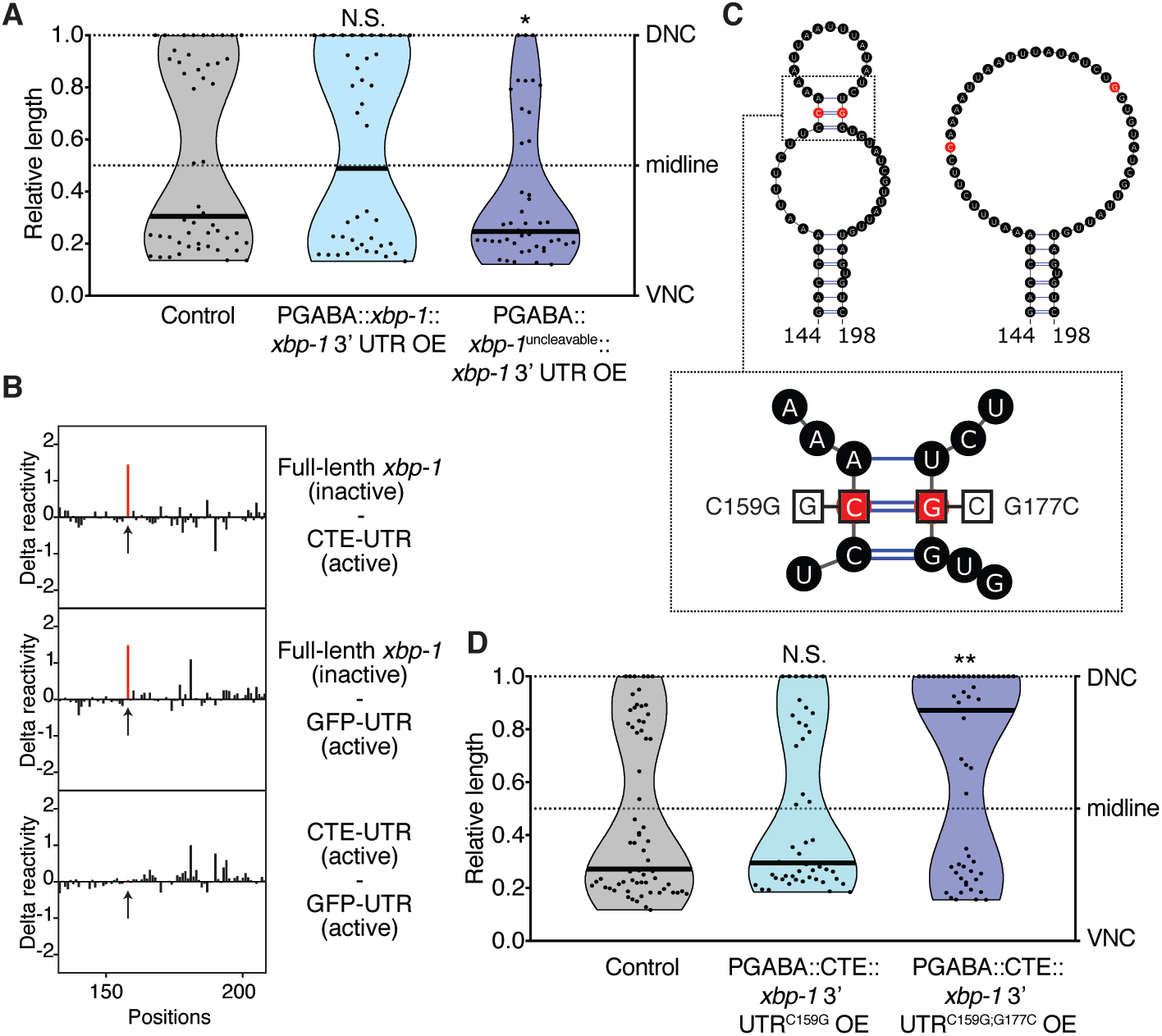
A single base pair within S^134-209^ is essential for *xbp-1* ncRNA function. (**A**) Overexpression of the full-length *xbp-1* transcript does not increase regeneration. *n* = 50, 41, and 45 from left to right. (**B**) Base-by-base comparisons of SHAPE reactivity difference across the length of S^134-209^. Arrows point to C159. (**C**) Predicted secondary structures of partial S^134-209^ based on SHAPE-MaP results. Left, the active forms (*xbp-1* 3’ fragment, CTE-UTR, and GFP-UTR). Right, the inactive form (full-length *xbp-1*). Box insert, diagram of point mutations used in (E) and (F). (**D**) Base pairing at C159 is essential for the overexpressed *CTE::xbp-1* 3’ UTR to increase regeneration. *n* = 67, 48 and 55 from left to right. In (A) and (D), black bar represents the median. N.S., not significant, \**P*<0.05, \*\**P*<0.01, 2-tailed K-S test.

We used SHAPE-MaP to compare the RNA structures of the active RNAs (*xbp-1* 3’ fragment, CTE-UTR, GFP-UTR) with the inactive RNA (full-length *xbp-1*). We focused on the S^134-209^ sequence, since our analysis demonstrated that it is necessary for function (Fig. 4D). SHAPE-MaP identified a single nucleotide within S^134-209^ whose reactivity differences correlated with the different function of the RNAs (Fig. 5B and S3A). Specifically, C159 shows low SHAPE reactivity and is predicted to be paired in a stem structure in the three active RNAs (Fig. 5C, left, and Fig. S3B). On the other hand, in the inactive full-length *xbp-1* RNA, C159 is highly accessible, indicating that the stem containing it does not form (Fig. 5C, right, and Fig. S3B). These *in vitro* SHAPE data suggest that pairing at C159 may be critical *in vivo* for the function of the *xbp-1* ncRNA. To test the functional importance of C159 pairing, we mutated it to G (C159G, Fig. 5C, box) in the CTE-UTR construct. This single mutation completely abolished the ability of the CTE-UTR construct to promote regeneration (compare Fig. 5D and 3C). Next, we introduced a compensatory mutation (G177C, Fig. 5C, box) that our structural analysis predicted would restore base pairing and stem formation. This second mutation restored the regeneration-promoting activity of the CTE-UTR construct (Fig. 5D). Together, these results confirm that the CTE-UTR construct acts as a ncRNA, and demonstrate that pairing at C159 is required for function.

### The endogenous *xbp-1* transcript has coding/non-coding dual output

The endogenous *xbp-1* transcript encodes the XBP-1 protein and has a well-established role in the XBP-UPR. Our experiments show that overexpressed fragments of this transcript can also act as ncRNAs. But does the endogenous *xbp-1* transcript give rise to a ncRNA in addition to its coding function? To determine the potential ncRNA function of the *xbp-1* 3’ fragment in its natural context, we used CRISPR to engineer the C159G (Fig. 5C, box) equivalent change into the endogenous *xbp-1* locus (*xbp-1(no-stem)*). In contrast to the *xbp-1(uncleavable)* and *xbp-1(deletion)* alleles, *xbp-1(no-stem)* animals have a normal XBP-1 protein-mediated UPR (Fig. 6A). Thus, the *xbp-1(no-stem)* allele does not affect the coding output of the endogenous *xbp-1* transcript, as is expected from a single nucleotide change in the 3’ UTR. However, *xbp-1(no-stem)* animals have significantly impaired regeneration compared to control animals (Fig. 6B). The loss of regeneration is nearly as strong as in the *xbp-1(uncleavable)* and *xbp-1(deletion)* alleles (Fig. 1F and 2D), indicating that C159 is required for the function of the endogenous, non-coding *xbp-1* 3’ fragment *in vivo*.

**Fig. 6.**
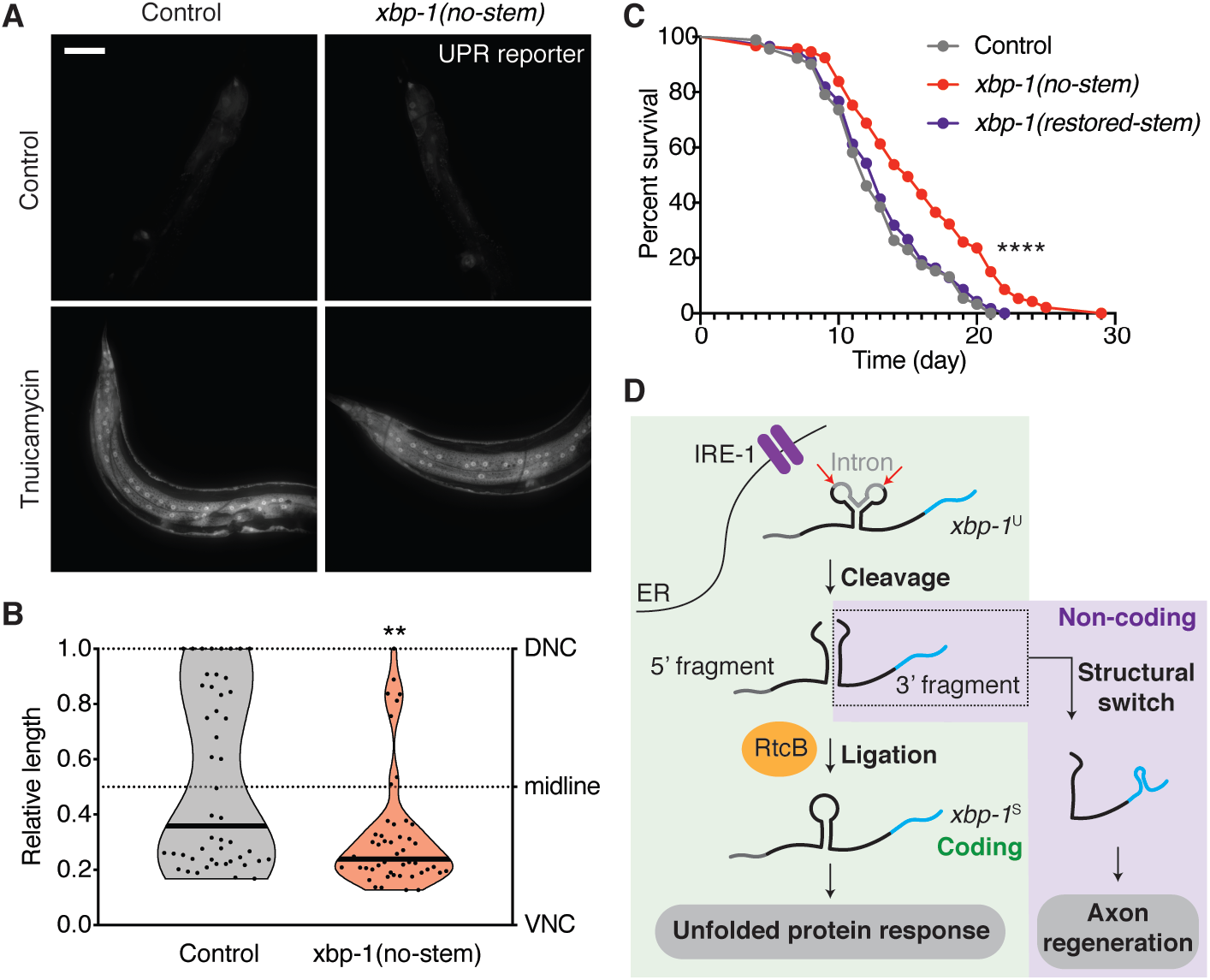
The endogenous *xbp-1* transcript has coding/non-coding dual output. (**A**) C159G mutation in the endogenous *xbp-1* locus does not affect the UPR. Tunicamycin treatment was at 5 μg/ml for 10hr. Scale bar, 50μm. (**B**) C159G mutation at the endogenous *xbp-1* locus decreases regeneration. *n* = 49 and 50 from left to right. Black bar represents the median. N.S., not significant, \*\**P*<0.01, 2-tailed K-S test. (**C**) The endogenous *xbp-1* ncRNA impacts animal life span. *n* = 91, 93 and 116 from top to bottom. \*\*\*\**P*<0.0001, log rank Mantel-Cox test. (**D**) Model of the coding and non-coding outputs of the *xbp-1* locus.

The non-coding output of the endogenous *xbp-1* transcript impacts more than just axon regeneration. We found that *xbp-1(no-stem)* animals have a significantly longer life span compared to control animals (Fig. 6C). Similar to regeneration, this effect is also dependent on the pairing at the C159 site, because the long life span was fully restored back to normal in *xbp-1(restored-stem)* animals, where we CRISPR engineered the G177C (Fig. 5C, box) equivalent change into the *xbp-1* locus to restore the base pairing of endogenous *xbp-1* ncRNA. Taken together, our data indicate that in its natural context, *xbp-1* mediates a non-coding pathway that is separable from its coding-dependent function; and that the endogenous *xbp-1* ncRNA affects both axon regeneration and animal life span (Fig. 6D).

## Discussion

### RNA processing intermediates with unidentified functions

The *xbp-1*^U^ transcript undergoes a well-conserved non-canonical splicing pathway, where an endonuclease and a ligase cooperate to generate the protein-coding *xbp-1*^S^ transcript. While previous studies of the output of the *xbp-1* pathway focused on the product of the *xbp-1*^S^ transcript, which encodes the critical UPR mediator XBP-1, our work revealed an unexpected role of a processing intermediate fragment of this pathway. We report that this intermediate RNA fragment functions as a ncRNA, entirely independently of the XBP-UPR.

More broadly, our data support the concept that RNA processing events like RNA cleavage can produce functional intermediate fragments in addition to the final product. This phenomenon has been observed previously, the most well-documented example being tRNA fragments (Hanada et al., 2013; Sobala and Hutvagner, 2011; Thompson and Parker, 2009). However, prior to this study, it was not known whether mRNA cleavage would also result in RNA fragment with non-coding functions. mRNA cleavage is a widespread cellular event. For example, during the regulated IRE1-dependent decay of mRNA (RIDD), numerous mRNA are cleaved by the endonuclease IRE-1 (Hollien et al., 2009). Our study therefore opens up new avenues for the generation of novel, functional ncRNAs.

### Roles of *xbp-1* in regulating the neuronal injury response

The *xbp-1* locus has been shown to function in neuronal injury across species (Hu et al., 2012; Kosmaczewski et al., 2015; Ohtake et al., 2018; Oñate et al., 2016; Song et al., 2015; Ying et al., 2015). It has been linked to axon regeneration itself, as well as to other cellular activities such as cell survival, myelin removal, and microphage infiltration (Hu et al., 2012; Oñate et al., 2016). Our work identifies a novel mechanism by which the *xbp-1* locus functions in neuronal injury: via a ncRNA generated by *xbp-1* mRNA cleavage.

Our finding deepens our understanding of how *xbp-1* mediates the neuronal injury response. Specifically, by cleaving and ligating the *xbp-1* mRNA, neurons can activate the XBP-UPR; and by cleaving but not ligating the *xbp-1* mRNA, neurons can activate a non-coding pathway mediated by the *xbp-1* 3’ fragment (Fig. 6D). The two pathways could be partially or preferentially activated to fine-tune the response to injury and other stresses via different cellular responses. Surprisingly, the *xbp-1* non-coding pathway described here is the major output of the *xbp-1* locus for activating axon regeneration and outgrowth after *C. elegans* axon injury, while the coding pathway has a relatively minor role. Further, the effect size of the *xbp-1* non-coding pathway on axon regeneration is very large, similar to the effect size of the well-characterized DLK-1 pathway (Hammarlund et al., 2009). Because the *xbp-1* non-coding pathway functions independently of the XBP-UPR, and can be separated from the other functions of RtcB, this pathway represents a powerful way to tune the response to injury without affecting these other basic cellular mechanisms.

### Regulation of the coding/non-coding dual outputs

RNA molecules are generally divided into two worlds with little overlap, coding and noncoding. Our data identify *xbp-1* as encoding the first ncRNA that is directly derived from the cleavage of a mRNA. We further show that the endogenous *xbp-1* transcript has coding/non-coding dual outputs. The coding output mediates the XBP-UPR. This is triggered by the buildup of unfolded proteins within the ER and involves expression of the chaperone BiP. The non-coding output is mediated by the *xbp-1* 3’ fragment. It impacts both axon regeneration and animal life span. Our work thus places the *xbp-1* RNA into the small group of RNAs that act both as coding and non-coding RNAs, sometimes called “coding and non-coding RNAs” (cncRNAs)(Crerar et al., 2019; DeJesus-Hernandez et al., 2011; Ji et al., 2015; Kumari, 2015; Mori et al., 2013; Sampath and Ephrussi, 2016). Such RNAs blur the line between the coding and non-coding RNA worlds and raise the question of how their dual roles are regulated.

In the case of *xbp-1*, the coding/non-coding dual outputs can be regulated by regulating cleavage and ligation. While regulation of cleavage of *xbp-1* mRNA by IRE-1 has been extensively characterized, far less is known about regulation of ligation by RtcB. However, multiple potential avenues for regulation exist. First, although *in vitro* RtcB exhibits RNA ligase activity on its own, *in vivo* it is thought to act as the catalytic subunit of a complex that includes archease and DDX1 (Desai et al., 2014, 2015; Popow et al., 2014). RtcB has also been reported to complex *in vivo* with multiple other accessory proteins like FAM98B and hCLE/C14orf166/RTRAF (Kanai et al., 2004; Pazo et al., 2019; Pérez-González et al., 2014). In addition, RNA ligation by RtcB requires specific chemistry at the RNA ends to be joined; a number of factors, including the cyclic phosphodiesterase CNP, the RNA cyclase RtcA (Unlu et al., 2018), as well as cytoplasmic capping machinery, could potentially alter this chemistry. It will be of interest to learn whether any of these potential modifiers are used as cellular mechanisms to regulate the balance between the coding and non-coding output of the *xbp-1* locus.

How do cleavage and ligation regulate the function of the *xbp-1* ncRNA? Our study indicates that it is through altering the structure of the ncRNA. Our data show that the function of the *xbp-1* ncRNA sequence depends on its molecular context. Although the entire ncRNA sequence is contained within the full-length transcript, it has no function in this context. Suppression of ncRNA function is likely due to interaction with specific sequences within the rest of the *xbp-1* transcript, since the *xbp-1* ncRNA is functional when it is placed in the context of a different mRNA (encoding GFP). We find that cleavage causes a structural switch in the *xbp-1* 3’ fragment, activating its non-coding function. Specifically, our analyses indicate that in the active *xbp-1* ncRNA, a critical base pair at the C159 site is formed. The formation of a stem structure may allow the ncRNA to interact with RNA binding proteins, which may facilitate the stability and/or function of the RNA. The cleavage-induced structural switch alters both the conformation and the function of the *xbp-1* RNA, and thus is a novel mechanism to switch between the coding and non-coding outputs of a cncRNA depending on the status of the cell.

## Acknowledgments

We thank WormBase and the Caenorhabditis Genetics Center (CGC), which is funded by the National Institute of Health (NIH) Office of Research Infrastructure Programs (P40 OD010440).

## Author contributions

X.L., J.-D.B., S.G.K, C.A.D., and M.H. designed experiments. X.L. performed the experiments and data analysis. S.G.K, B.I.M, J.-D.B. performed the RNA-seq experiments. J.-D.B. performed the splice junction analysis with the help of S.G.K and X.L.. J.-D.B. performed the SHAPE-MaP experiments and processed the SHAPE-MaP data. C.A.D performed the life span assay. All authors discussed and interpreted the results. M.H. supervised the project, with the contribution of A.J.G.. X.L. and M.H. prepared the manuscript with input from the other authors.

## Data availability

All data are available in the main text or the supplementary materials.

## Supplementary Materials

**Fig. S1.**
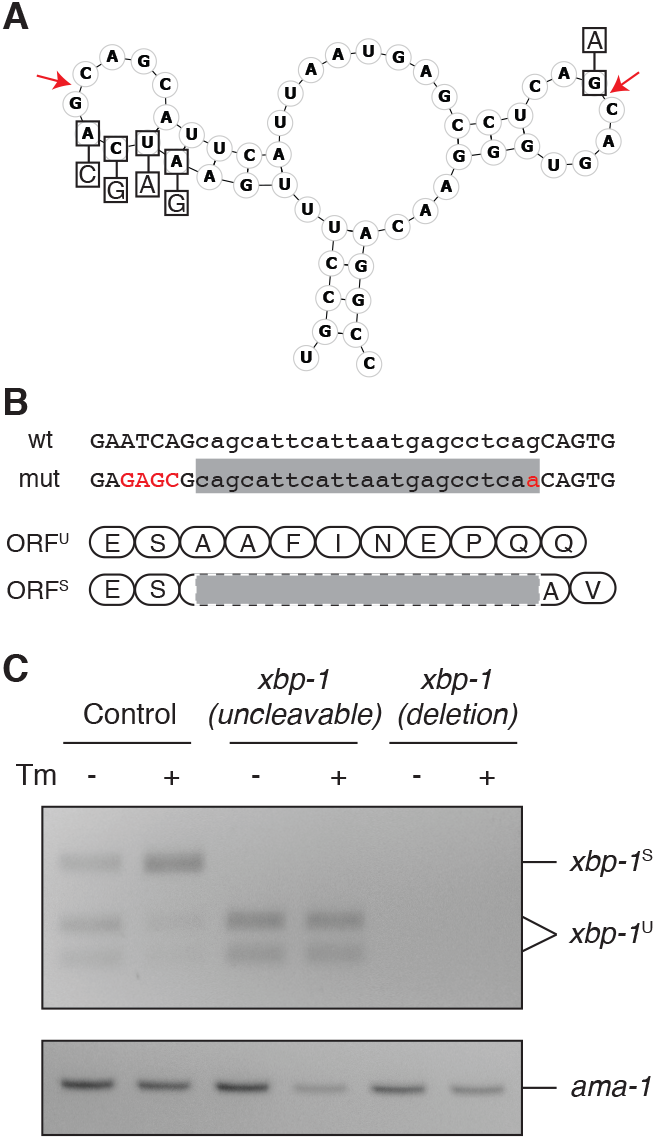
Generation of an uncleavable allele of *xbp-1*. (**A**) Diagram of the uncleavable *xbp-1* mutation. Arrows point to the two cleavage sites. (**B**) DNA sequence surrounding the alterations in the uncleavable allele (mut). Changed bases are shown in red. Lowercase indicates the sequence removed by non-canonical splicing. The translations below (in the two different frames) are not affected. (**C**) RT-PCR amplifying both forms of *xbp-1* transcript from RNA extracted from animals with indicated genotypes. The amplification products were digested with a restriction enzyme that specifically cuts *xbp-1*^U^ before visualization by gel. Tm, tunicamycin.

**Fig. S2.**
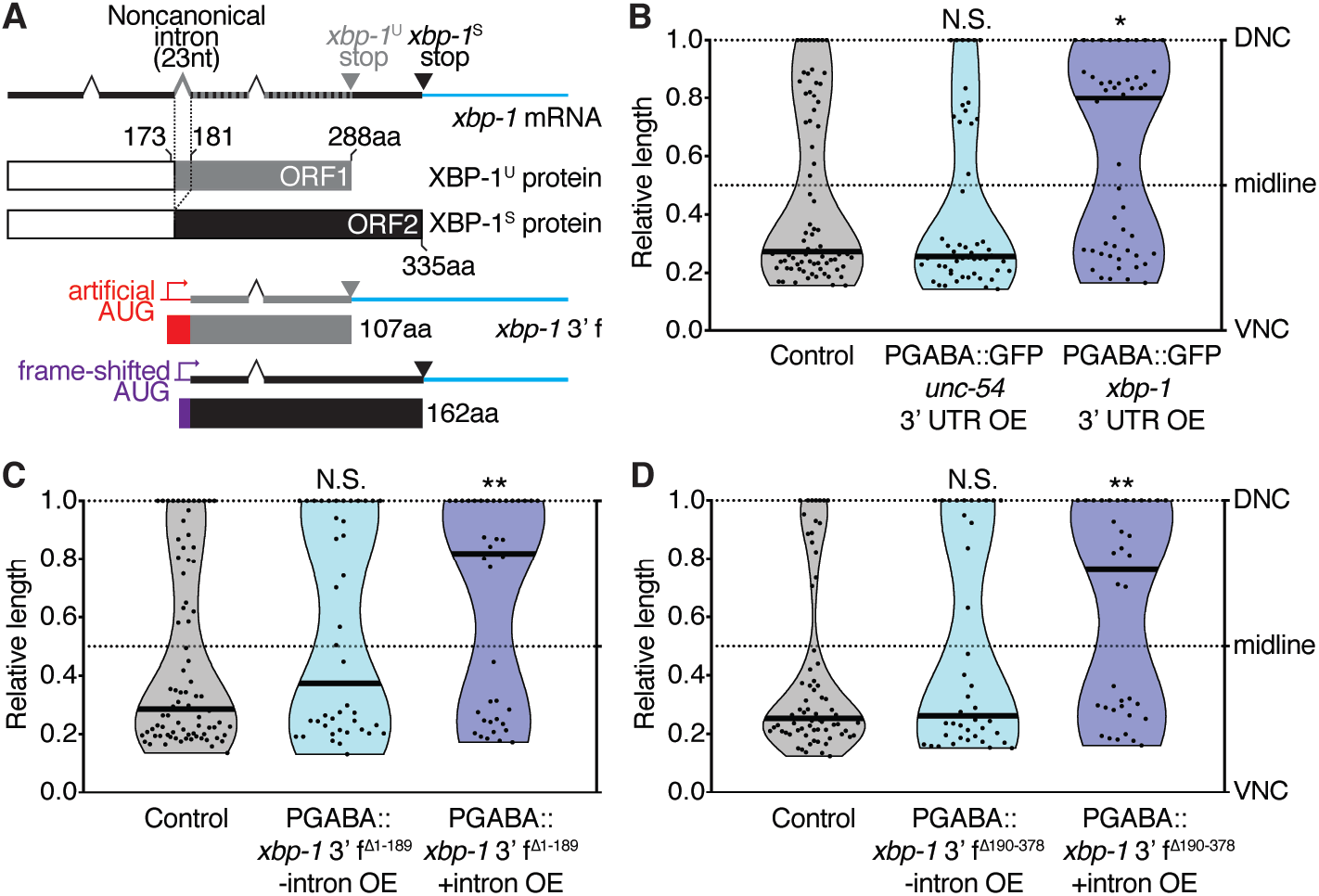
The function of the *xbp-1* 3’ UTR is independent of the *xbp-1* coding sequence but requires an intron. **(A)** Diagram of the *xbp-1* mRNA and the encoded proteins. Non-canonical splicing of the 23nt intron causes a frameshift. The two different open reading frames (ORFs) are indicated with different colors. **(B)** Overexpression of GFP increases regeneration only when fused with the *xbp-1* 3’ UTR. *n* = 76, 53 and 50 from left to right. (**C** and **D**) The *xbp-1* 3’ fragment increases regeneration only in the presence of an intron. Deletion constructs Δ1-189 (C, *n* = 82, 42, and 39) and Δ190-378 (D, *n* = 71, 40, and 36) shown in Fig. 3D with or without an intron were tested. The with-intron data are the same as in Fig. 3E. In (B-D), black bar represents the median. N.S., not significant, \**P*<0.05, \*\**P*<0.01, 2-tailed K-S test.

**Fig. S3.**
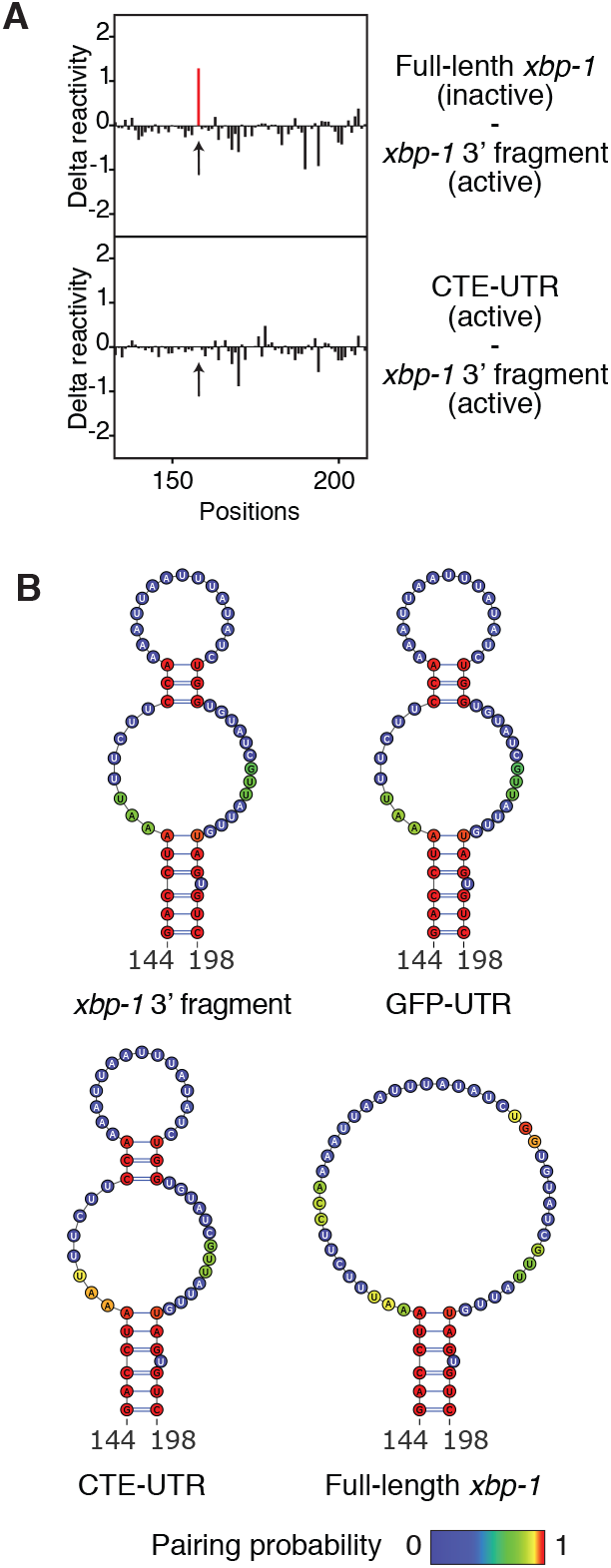
SHAPE-MaP reveals structural difference between *xbp-1* 3’ UTRs that are active and inactive in promoting axon regeneration. (**A**) Base-by-base comparisons of SHAPE reactivity difference across the length of S^134-209^. Arrows point to C159. (**B**) Predicted secondary structures of partial S^134-209^ in the indicated RNA molecules based on SHAPE-MaP results.

### Materials and Methods

#### C.elegans

*C. elegans* were maintained on nematode growth media (NGM) plates seeded with *Escherichia coli* OP50 at 20 °C. The strains used are listed below. For transgenic animals with extrachromosomal arrays, the relevant constructs as indicated below along with 2.5ng/μl *Pmyo-2*::mCherry (injection marker) and DNA ladder (Promega, G5711, to adjust the total concentration of the injection mix), were microinjected into the gonads of young adults. Transgenic animals were first selected based on the expression of *Pmyo-2*::mCherry, and then confirmed by PCR genotyping of the relevant constructs.

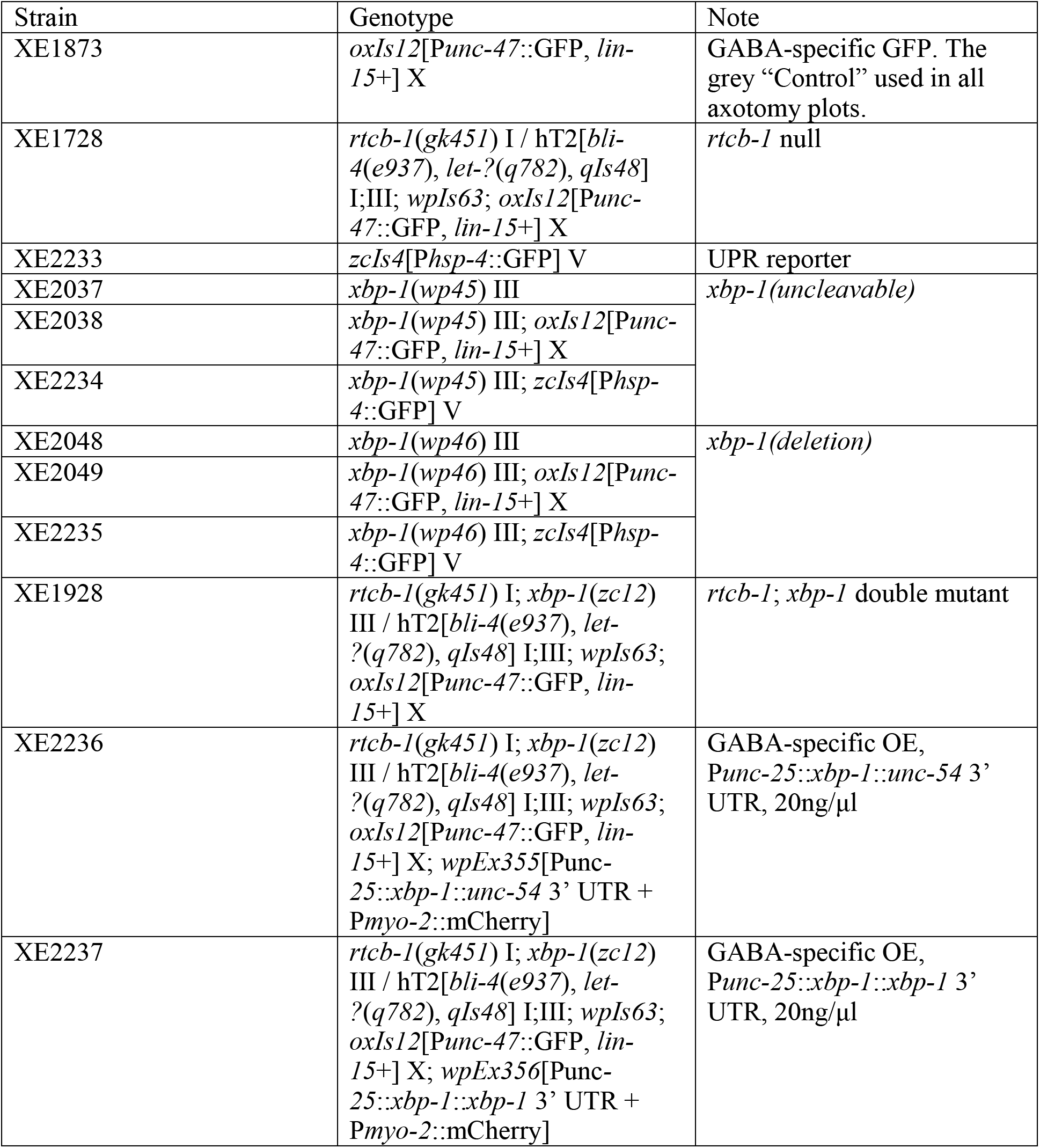
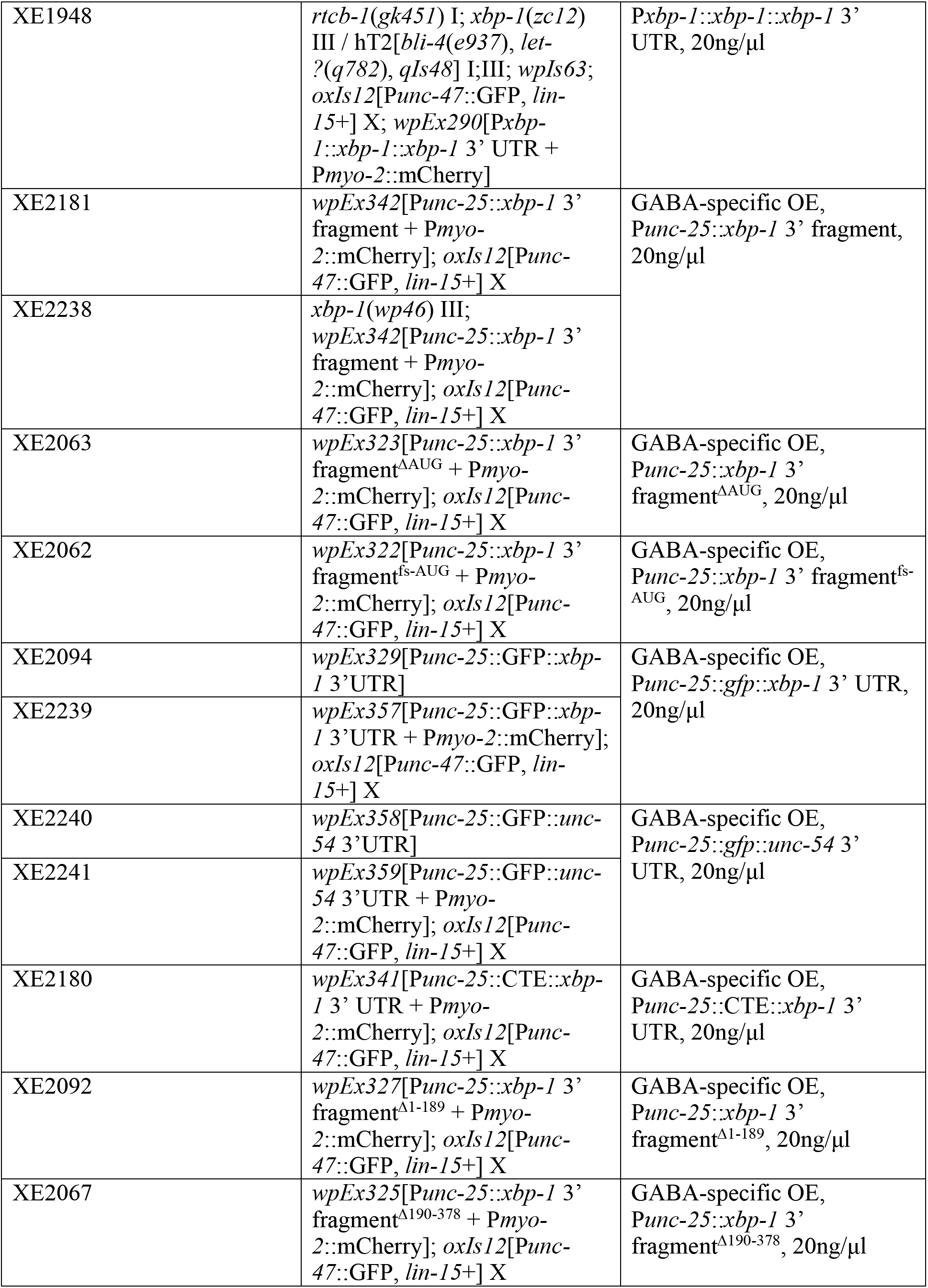
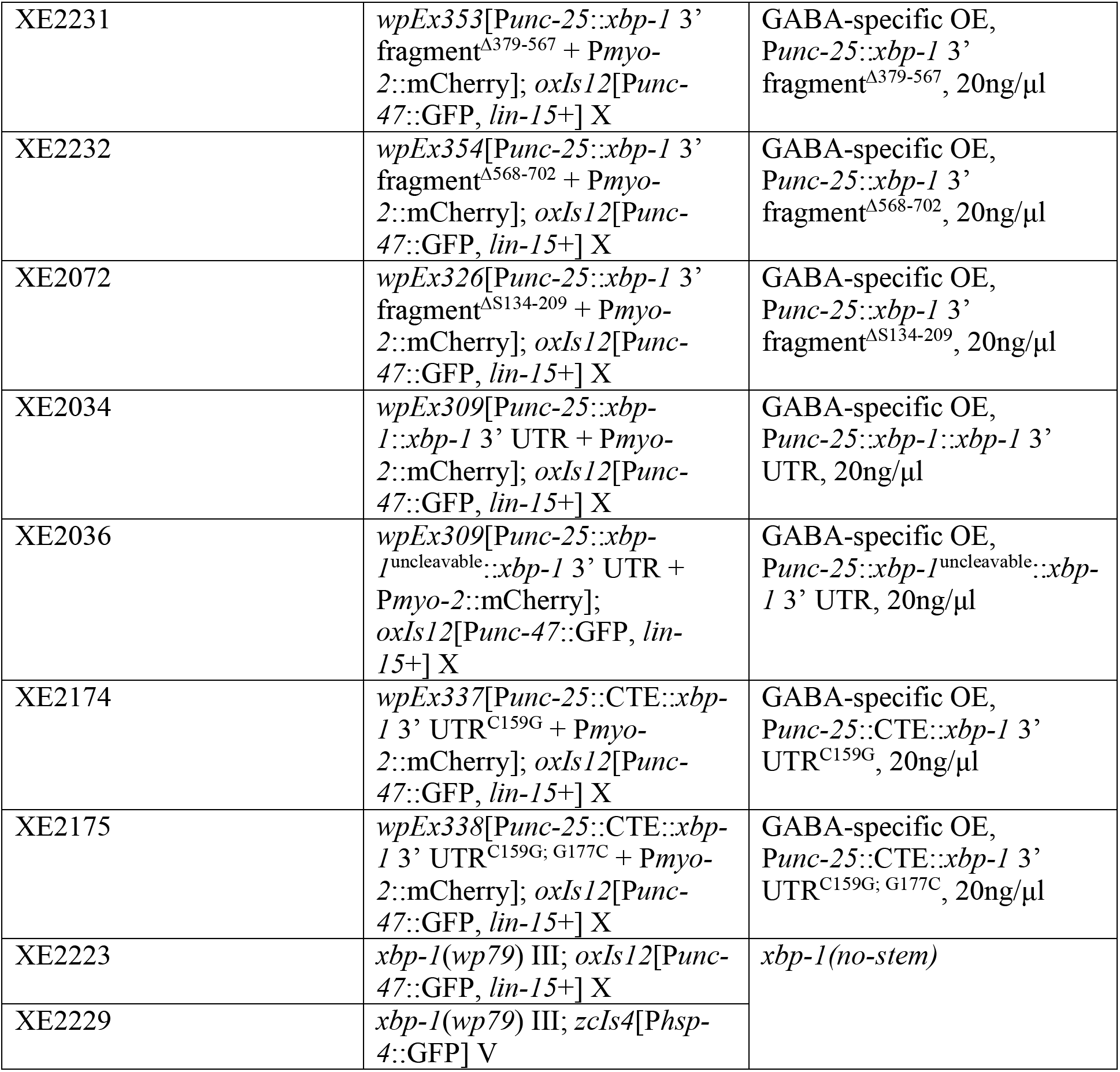

#### CRISPR Mutations

*The four xbp-1 alleles (wp45, wp46, wp79, wp80)* were generated by CRISPR as previously described(Arribere et al., 2014; Farboud and Meyer, 2015; Paix et al., 2014). The obtained mutants were confirmed by sequencing and were outcrossed >3X prior to any experiments.

#### Molecular Biology

Gateway recombination (Invitrogen) was used to generate *Punc-25::xbp-1::unc-54 3’ UTR, Punc-25::xbp-1::xbp-1 3’ UTR, Pxbp-1::xbp-1::xbp-1 3’ UTR, Punc-25::gfp::xbp-1 3’ UTR, and Punc-25::gfp::unc-54 3’ UTR. Gibson cloning was used to generate Punc-25::CTE::xbp-1* 3’UTR and *Punc-25::xbp-1* 3’ fragment. The CTE sequence was synthesized by IDT. Other entry pieces were amplified using Phusion polymerase (NEB, M0530) from worm lysate or previous plasmids (Kosmaczewski et al., 2015). Gibson assembly was also used to generate the various deletion constructs on the basis of the *Punc-25::xbp-1* 3’ fragment construct. Site-directed mutagenesis was used to introduce the C159G and G177C point mutations to the *Punc-25::CTE::xbp-1* 3’UTR construct.

All construct and primer sequences are available upon request.

#### *xbp-1* RT-PCR assay

Worms of N2 wild-type, *xbp-1(wp45)* or *xbp-1(wp46)* were harvested from fully-populated NGM plates just before starving. Next they were washed multiple times in M9 buffer, nutated in M9 with or without 5μg/ml tunicamycin for 3h, spun down, resuspended in TRIzol reagent (Invitrogen), frozen at −80°C overnight, and then subjected to freeze/thaw in liquid nitrogen/37°C incubator 3 times. RNA was isolated from the aqueous phase following manufacturer’s instructions. First-strand cDNA was synthesized with AffinityScript Multiple Temperature cDNA Synthesis Kit (Agilent, #200436). PCR was carried out using Phusion polymerase (NEB, M0530) with the following intron-spanning primers: Forward GAGACAAAAAGAAG-GAAAGATCAGC and Reverse CTCCGCTTGGGCT-CTTGAGATG for *xbp-1*^U^ and *xbp-1^S^;* Forward TGTCAGGATCGAAGGGATCGAAG and Reverse CGGTGAGGTCCAT-TCTGAAATC for *ama-1* control. Products were digested with MseI (NEB, R0525) and then resolved on a 2% agarose gel.

#### UPR Fluorescent Reporter Assay

UPR assay was performed as described previously (Kosmaczewski et al., 2015). Animals at the L4 stage were placed on NGM plates with or without 5 μg/ml tunicamycin for 10-24h as indicated. P*hsp-4*::GFP expression was then visualized by imaging.

#### Imaging of live animals

Animals were mounted on 3% (wt/vol) agarose pads and immobilized using 0.1% Levimisole (Santa Cruz, sc-205730). Images were taken with Olympus BX61 and processed with Micro-Manager.

#### Laser Axotomy

Laser axotomy was performed as described previously (Kosmaczewski et al., 2015). Briefly, worms at L4 stage were mounted on a 3% (wt/vol in M9 buffer) agarose pad, immobilized by 0.05μm microbeads (Polysciences, #08691) with 0.02% SDS, and visualized with a Nikon Eclipse 80i Microscope. 3-4 of the 7 most posterior ventral and dorsal D-type (VD/DD) GABA motor neurons were severed using a Photonic Instruments Micropoint Laser at 10 pulses and 20 Hz. Worms were recovered to seeded NGM plates and scored for axon regeneration 24h (unless otherwise stated) after axotomy. The relative lengths (Fig. 1B) of all successfully cut axons (indicated by the presence of a severed distal stump) were measured with ImageJ and plotted with GraphPad Prism. Two-tailed *P* values were calculated using the Kolmogorov-Smirnov test (2-tailed K-S test). All scoring processes were carried out blindly. Measurements were taken from distinct samples for each plot shown.

#### Northern blots

~2,000 L4 N2 wild-type or *rtcb-1(gk451)* worms were picked to NGM plates with or without 50ug/ml tunicamycin (*rtcb-1* homozygotes were isolated from balanced heterozygotes), incubated at 20°C for 3h, and harvested from the plates with M9 buffer. RNA was isolated as in the *xbp-1* RT-PCR assay. 10μg of total RNA was separated in 1.2% agarose-37% formaldehyde gels and transferred to Zeta-Probe membranes (Bio-Rad, 1620165) by capillary transfer in 20X SSC buffer for 22h. The blots were first hybridized and visualized with the following [^32^P]-labeled oligonucleotide probes against the *xbp-1* sequence 3’ to the IRE-1 cleavage sites (*xbp-1* probes), and then stripped, and hybridized and visualized with *act-1* probes as control. The hybridized signals were visualized with the Storm 860 Molecular Imager (GMI).

*xbp-1* probes: 1. TGATGTTGTTGTTGATGGAGGTGGATCGGGCCTG; 2. GTGTCCAT CTTCTTGTTGCGATCCATGTGGTTG; 3. CCTTCCTCAATGTAGCCAGCAACGAATCGA TC; 4. GACGGGGACATGGCTAGGGAATTCGGCGAAAGTG; 5. GTTCCACATCCCAAAA GCTCATCATCCCAGTC; 6. GGTAAGCAGCTCGTCGGTTCCAGTTCCAGTTTC; 7. GCTT GGGCTCTTGAGATGTTCGAGGATTTGTTCG.

*act-1* probes: 1. GTAAGGATACCTCTCTTGGATTGGGCCTCGTCTC; 2. CTTCATGGT TGATGGGGCAAGAGCGGTGATTTC.

#### RNA-seq and splice junction analysis

RNA-seq experiments were performed to identify RtcB-dependent changes in mRNA splicing. Samples were prepared in triplicate from the following four groups: control, *rtcb-1(gk451), unc-70(s1502)*, and *rtcb-1(gk451); unc-70(s1502)*. For each replicate, ~2,000 L4 worms were gathered. Total RNA was extracted using TRIzol as described above. Poly-A+ RNAs were isolated using oligo d(T)25 magnetic beads (New England BioLabs) following the manufacturer’s protocol and eluted in 11 μL of water. Reverse transcription was performed in 20 μL at 25 °C for 10 min, 42 °C for 30 min, 50 °C for 10 min, 55 °C for 20 min, and 60 °C for 20 min, using Superscript III (Invitrogen) and 5’-AGACGTGTGCTCTTCCGATCTNNNNNN-3’ (IDT DNA) reverse transcription random primer. The reaction was then heat inactivated at 75 °C for 15 min and RNAse H treated at 37 °C for 15 min. cDNA samples were purified using 36 μL of AMPure XP beads (Beckman Coulter Life Sciences) following the manufacturer’s protocol and eluted in 10 μL of water. Purified cDNA samples were ligated to a ssDNA adaptor (/5Phos/NNNNNGATCGTCGGACTGTAGAACTCTGAAC/3InvdT/) at their 3’-end using CircLigase ssDNA Ligase (Epicentre) following the manufacturer’s protocol with the addition of 1M betaine and 10% PEG 6000. Ligation reactions were incubated at 60 °C for 2h, at 68 °C for 1h and heat inactivated at 80 °C for 10 min. 10 μL of water was added to each reaction and ligated products were purified using 36 μL of AMPure XP beads (Beckman Coulter Life Sciences) and dissolved in 16 μL of water. Purified ligated cDNA samples were then PCR amplified using Illumina sequencing adapters, keeping the number of cycles to the minimum needed for the detection of amplified products (8-12 cycles), and gel purified on 2% agarose gel to remove adaptor-adaptor dimers. Purified libraries were sequenced on Illumina HiSeq 2000/2500 machines producing single-end 76 nucleotide reads.

Following the library preparation protocol, raw reads contained the following features: NNNNN-insert-adapter, where the 5N sequence composes the Unique Molecular Identifier (UMI), and adapter is the 3’-Illumina adapter that can occasionally be present within the read (mainly in adapter-adapter dimers). The UMI was used to discard PCR duplicates and count single ligation events. Base calling was performed using CASAVA-1.8.2. The Illumina TruSeq index adapter sequence was then trimmed when present by aligning its sequence, requiring a 100% match of the first five base pairs and a minimum global alignment score of 60 (Matches: 5, Mismatches: −4, Gap opening: −7, Gap extension: −7, Cost-free ends gaps). The UMI was clipped from the 5’-end and kept within the read name, for marking PCR duplicates. Reads were then depleted of rRNA, tRNA, snRNA, snoRNA and miscRNA, using Ensembl 80 annotations, as well as from RepeatMasker annotations, using strand-specific alignment with Bowtie2 v2.2.4 (Langmead and Salzberg, 2012). Next, reads were aligned to the *C. elegans* WBcel235 genome assembly using STAR version 2.4.2a (Dobin et al., 2013) with the following non-default parameters: *--alignEndsType EndToEnd --outFilterMultimapNmax 100 --seedSearchStartLmax 15 --sjdbScore 10 --outSAMattributes All −limitBAMsortRAM 3221225472*. Genomic sequence indices for STAR were built including exon-junction coordinates from Ensembl 80. Finally, the STAR SJ.out.tab output files were parsed and the coverage of individual splice sites was compared between control and *rtcb-1(gk451)* (Table S1, “no neuronal injury” tab), and between *unc-70(s1502)* and *rtcb-1(gk451); unc-70(s1502)* (Table S1, “neuronal injury” tab). For each comparison, we considered only splice sites i) were detected in at least two out three replicates of genotype 1, ii) were supported by at least three uniquely mapped reads in genotype 1 and iii) had a splice site coverage difference of at least 2-fold between the two genotypes after normalizing for changes in RNA abundance (equation 1). A value of 0.01 was added to splice site coverages equal to 0 in genotype 2. Only uniquely mapped reads were considered in the calculation of splice site coverages and RPKM (Read Per Kilobase Million) values. Non-canonical splice site with >10-fold difference in either of the two comparisons were annotated based on WormBase annotation (Table S2). Venn diagrams were drawn with Meta-chart.

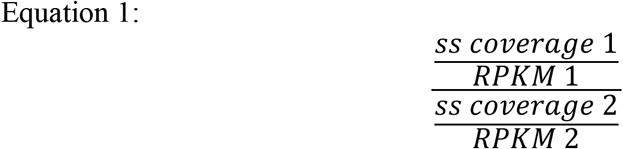

where *ss coverage 1* is the splice site coverage in genotype 1, *RPKM1* is the mRNA’s RPKM value in genotype 1, *ss coverage 2* is the splice site coverage in genotype 2, and *RPKM2* is the mRNA’s RPKM value in genotype 2.

#### SHAPE-MaP

RNA species were transcribed *in vitro* from PCR products using the AmpliScribe T7-Flash transcription kit (Epicentre), and purified using RNeasy columns (QIAGEN). The Reverse primer CAAAAATGTATGGCTGTTTTTATTAAATATACAAATGGGGGAATTG and the indicated Forward primers were used to generate the following transcripts:

1. *xbp-1* 3’ fragment TAATACGACTCACTATAGGCAGTGGGAACAGGCCCGATC cagtgggaacaggcccgatccacctccatcaacaacaacatcagcaaccaactccgtcgtatggattccaagaagaacaacac aatcagtgtggatatgtatctaactatcatctcgattctatgcaaccacatggatcgcaacaagaagatggacacctcgaacaaatc ctcgaacatctcaagagcccaagcggagagttcgatcgattcgttgctggctacattgaggaaggagcagacggttatgcagcg tcttgttcaagcggatccatgtacacatcttcagaaacgcgtgaaacactttcgccgaattccctagccatgtccccgtcgatgagc agctcgagcactgactgggatgatgagcttttgggatgtggaaccgaaactggaactggaaccgacgagctgcttaccgaccc cggaaactggaactttgaaactttcgacgaaaattcaatcgacctaaatttcttccaaaattaatttatatctggtgtatcgttattgtag tgtcctagtgaatttccctcgttattattttctccccaaaaatctctccaaatttctccatttcctctctggtattttcttcttttcctcttttcctt ttgtacaaccccaactttcgtattattatttacccttattaaatttccctcccccaaagctcattgattccaattcccccatttgtatatttaa taaaaacagccatacatttttg
2. *gfp::xbp-1* 3’ UTR (GFP-UTR) TAATACGACTCACTATAGGATGAGTAAAGGAGAAGAACTTTTCACTGGAGatgagtaaaggagaagaacttttcactggagttgtcccaattcttgttgaattagatggtgatgttaatgggcacaaattttctgtcagtggagagggtgaaggtgatgcaacatacggaaaacttacccttaaatttatttgcactactggaaaactacctgttccatggccaacacttgtcactactttctgttatggtgttcaatgcttctcgagatacccagatcatatgaaacggcatgactttttcaagagtgccatgcccgaaggttatgtacaggaaagaactatatttttcaaagatgacgggaactacaagacacgtgctgaagtcaagtttgaaggtgatacccttgttaatagaatcgagttaaaaggtattgattttaaagaagatggaaacattcttggacacaaattggaatacaactataactcacacaatgtatacatcatggcagacaaacaaaagaatggaatcaaagttaacttcaaaattagacacaacattgaagatggaagcgttcaactagcagaccattatcaacaaaatactccaattggcgatggccctgtccttttaccagacaaccattacctgtccacacaatctgccctttcgaaagatcccaacgaaaagagagaccacatggtccttcttgagtttgtaacagctgctgggattacacatggcatggatgaactatacaaatagccatgtccccgtcgatgagcagctcgagcactgactgggatgatgagcttttgggatgtggaaccgaaactggaactggaaccgacgagctgcttaccgaccccggaaactggaactttgaaactttcgacgaaaattcaatcgacctaaatttcttccaaaattaatttatatctggtgtatcgttattgtagtgtcctagtgaatttccctcgttattattttctccccaaaaatctctccaaatttctccatttcctctctggtattttcttcttttcctcttttccttttgtacaaccccaactttcgtattattatttacccttattaaatttccctcccccaaagctcattgattccaattcccccatttgtatatttaataaaaacagccatacatttttg
3. CTE: *:xbp-1* 3’ UTR (CTE-UTR) TAATACGACTCACTATAGGTCCCCTGTGAGCTAGACTGGAC tcccctgtgagctagactggacagccaatgacgggtaagagagtgacatttctcactaacctaagacaggagggccgtcaaagctactgcctaatccaatgacgggtaatagtgacaagaaatgtatcactccaacctaagacaggcgcagcctccgagggatgtgtccatgtccccgtcgatgagcagctcgagcactgactgggatgatgagcttttgggatgtggaaccgaaactggaactggaaccgacgagctgcttaccgaccccggaaactggaactttgaaactttcgacgaaaattcaatcgacctaaatttcttccaaaattaatttatatctggtgtatcgttattgtagtgtcctagtgaatttccctcgttattattttctccccaaaaatctctccaaatttctccatttcctctctggtattttcttcttttcctcttttccttttgtacaaccccaactttcgtattattatttacccttattaaatttccctcccccaaagctcattgattccaattcccccatttgtatatttaataaaaacagccatacatttttg
4. *xbp-1::xbp-1* 3’ UTR (full-length *xbp-1*) TAATACGACTCACTATAGGATGAGCAACTATCCAAAACGTATTTATG atgagcaactatccaaaacgtatttatgtgctcccagcacgccacgtggcagcgccacagcctcagagaatggctcccaagcgt gcacttccaacagaacaagttgtcgcacaacttcttggcgatgatatgggaccatctgggccacgcaaaagagaacgactgaat catttgagtcaggaggagaaaatggatcgtcggaaacttaaaaatcgagtcgcagcccaaaatgctagagacaaaaagaagga aagatcagcaaagatcgaggatgtgatgcgcgatctggtggaggagaaccgccggctccgcgctgaaaacgaacgtcttcgc cgtcaaaataaaaatcttatgaaccagcagaacgagtccgtcatgtatatggaagagaacaacgaaaacttgatgaacagcaat gatgcatgcatctaccagaacgtcgtctacgaagaagaagtcgtcggtgaggttgcaccagttgtcgtcgtcggaggagaggat cgccgtgcctttgaatcagcagcattcattaatgagcctcagcagtgggaacaggcccgatccacctccatcaacaacaacatca gcaaccaactccgtcgtatggattccaagaagaacaacacaatcagtgtggatatgtatctaactatcatctcgattctatgcaacc acatggatcgcaacaagaagatggacacctcgaacaaatcctcgaacatctcaagagcccaagcggagagttcgatcgattcg ttgctggctacattgaggaaggagcagacggttatgcagcgtcttgttcaagcggatccatgtacacatcttcagaaacgcgtga aacactttcgccgaattccctagccatgtccccgtcgatgagcagctcgagcactgactgggatgatgagcttttgggatgtgga accgaaactggaactggaaccgacgagctgcttaccgaccccggaaactggaactttgaaactttcgacgaaaattcaatcgac ctaaatttcttccaaaattaatttatatctggtgtatcgttattgtagtgtcctagtgaatttccctcgttattattttctccccaaaaatctct ccaaatttctccatttcctctctggtattttcttcttttcctcttttccttttgtacaaccccaactttcgtattattatttacccttattaaatttc cctcccccaaagctcattgattccaattcccccatttgtatatttaataaaaacagccatacatttttg

For each transcript, 1 μg of RNA in 14 μl final volume was heated to 95 °C for 2 min, placed on ice for 2 min, folded at 22 °C for 20 min with the addition of 4 μl of 5X folding buffer (500 mM tris HCl pH 7.5, 500 mM KCl and 50 mM MgCl2), probed at 22 °C for 5 min with the addition of 2 μL NAI (2 M, Sigma-Aldrich) or DMSO, and quenched by the addition of 90 μL of stop solution (3 M β-mercaptoethanol, 508 mM sodium acetate and 15 μg glycoblue). Following a 5 min incubation at room temperature, samples were ethanol precipitated, washed with 70% ethanol, resuspended in water, and subjected to MaP reverse transcription (requiring Superscript II and the addition of Mn^2+^ to the RT buffer(Smola et al., 2015)) using random nonamer primers. Double-stranded cDNA was synthesized in Second Strand Synthesis Reaction Buffer (NEB) with Second Strand Synthesis Enzyme mix (NEB), purified using PureLink PCR micro spin column (Thermo Fisher Scientific), eluted in water, and sent to the Yale Center for Genome Analysis to be fragmented using the Nextera DNA Flex Library Prep Kit (Illumina) and sequenced on an Illumina NovaSeq sequencer producing paired-end 150 nucleotide reads. Raw sequencing data were processed using the ShapeMapper pipeline (version 2.1.3)(Siegfried et al., 2014; Smola et al., 2015) with default parameters. SHAPE reactivities were only computed for nucleotides possessing sequencing depths above 1,000 in both modified and untreated samples. Nucleotides not passing this filter were treated as “no data” and excluded from downstream analysis. Secondary structure predictions and base pairing probabilities were generated using the SuperFold algorithm and SHAPE reactivities as restraints(Smola et al., 2015) with the following parameters: SHAPEintercept = −0.6, SHAPEslope = 1.8, trimInterior = 50, partitionWindowSize = 1200, PartitionStepSize = 100, foldWindowSize = 3000, foldStepSize = 300, maxPairingDist = 600.

#### Life span assay

For each genotype, ~100 L4 animals were picked to NGM plates seeded with OP50 bacteria. The animals were kept at 20°C and fed with OP50 during the assay. During the first 5 days, animals were transferred to a fresh seeded plate every other day to separate them from their off springs. Viability was scored every other day. Death was scored by failure to respond to touching. Survival curves were plotted with GraphPad Prism using the Kaplan Meier Method. Significance was calculated using the log rank Mantel-Cox test.

